# Optimized protocols for commonly-used murine models of heart failure with preserved ejection fraction

**DOI:** 10.1101/2025.01.22.634221

**Authors:** Bailey McIntosh, Ali Mohamed Elbassioni, Anmar Raheem, Eilidh McDonald, Stuart A. Nicklin, Yen Chin Koay, Ewan R. Cameron, Christopher M. Loughrey, John F. O’Sullivan

## Abstract

**Background:** HFpEF is a leading cause of death worldwide and clinically relevant preclinical models are required to identify new therapeutic targets. The most clinically representative murine models of heart failure with preserved ejection fraction (HFpEF) in common use include a “2-hit” model combining metabolic stress with hypertension (high-fat diet [HFD] + N(gamma)-nitro-L-arginine methyl ester [L-NAME]) and a “3-hit” model that includes age as an additional “hit” (age + HFD + deoxycorticosterone pivalate [DOCP]). However, both models have reproducibility challenges, and sub-strain and sex dependency. Here we optimize both preclinical models to overcome these challenges.

**Methods:** In this study we optimized both models: (1) The 2-hit model was optimised to reproduce HFpEF (defined as the induction and maintenance of obesity, hypertension, diastolic dysfunction, left ventricular hypertrophy, lung congestion, and exercise intolerance) in both C57BL/6N and 6J mice using increasing L-NAME doses (0.5 g/L to 1.75 g/L) and protocol lengths (7 weeks to 13 weeks); and (2) The 3-hit model used 12-week-old C57BL/6N and 6J mice and two aging protocols were compared: HFD for 7 months, or healthy chow for 5 months then high fat diet for 7 months. After HFD, mice received an intraperitoneal injection of DOCP to induce hypertension *via* sodium retention. To enhance and prolong the effect of DOCP, mice received 1% NaCl drinking water at the time of injection until sacrifice, henceforth called “4-hit”. To ensure the phenotype was maintained, a second bolus of DOCP was administered 8 weeks after the first.

**Results:** For the 2-hit protocol, HFpEF was successfully induced in C57BL/6J mice when exposed to a 13-week L-NAME protocol with gradually increasing dosage from 1.0 g/L to 1.75 g/L. C57BL/6N mice showed the desired parameters after 7-weeks of 0.5 g/L L-NAME, which were not augmented by increased dosage or time administered. For the 4-hit mice, after addition of 1% NaCl drinking water following DOCP administration, a clear HFpEF phenotype was observed in C57BL/6N and 6J mice in both male and females, and maintained for up to 12 weeks.

**Conclusions:** Our modifications ensure the 2-hit model is equally effective in both commonly used J and N substrains of C57BL/6 mice. Our 4-hit model overcomes the challenges of the 3-hit model, enhances reproducibility and robustness, which we demonstrate across sexes and substrains. Both of these new protocols will enhance clinically relevant mechanistic studies on HFpEF.

## Introduction

Heart failure (HF) is a clinical syndrome where the heart cannot pump blood at a rate commensurate with the body’s needs (Heart Failure with reduced Ejection Fraction, HFrEF) or only at the cost of increased filling pressures (Heart Failure with preserved Ejection Fraction, HFpEF) ^1^. HFpEF accounts for at least 50% of heart failure cases ^1,2^, presenting with clinical symptoms of HF such as exercise intolerance and fluid overload, frequently with associated atrial fibrillation, obesity, and metabolic syndrome ^3^.

While mice have proven to be effective models for human disease, replicating complex conditions such as HFpEF remains challenging. A commonly used murine model is the “2-hit” model developed by Schiattarella *et al*. ^4^ which combines a high-fat diet to perturb metabolism with the nitric oxide (NO) synthase inhibitor L-NAME to induce hypertension and nitrosative stress. Another, “3-hit”, model adds aging to hypertensive and metabolic stress ^5^ and has been described as amongst the most clinically representative models ^6,7^. In this approach, 3-month-old mice are fed a high-fat diet for 13 months to induce metabolic stress, and in the final month, desoxycorticosterone pivalate (DOCP) is administered intraperitoneally to induce hypertension and systemic inflammation.

Although both models have accelerated mechanistic research in HFpEF, several challenges and limitations remain. The 2-hit model was reported as being less reliable in female C57BL/6N mice ^8^. However, this protocol reproducibly produced the full spectrum of HFpEF in female C57BL/6J mice, as reported by us and others ^9–11^ . Others have shown that the 2-hit model is generally more successful in N than J strains, regardless of sex ^12^. Many genetically modified mouse models are generated in C57BL/6J strains and so it is important to be able to reproduce HFpEF in this strain. With regards to the “3-hit” model, we consistently encountered challenges in reproducibly generating HFpEF in mice. Therefore, in this manuscript, we sought to determine the reasons underpinning these inconsistencies and to optimize protocols for improved reproducibility and clinical relevance.

## Methods

### Animal husbandry and HFpEF models

Animal experiments were performed under the approval of University of Glasgow Animal Welfare and Ethical Review Body, Glasgow, UK (project license no. P05FEIF82 and subsequently PP4465428) and the Animal Ethics Committee of The University of Sydney (Project 2023/2274), Sydney, Australia. Animals were housed in controlled environments with a 12-hour light/dark cycle and had access to food and water *ad libitum*.

For the 2-hit protocol, 10-12 week C57BL/6J male mice were purchased from Envigo, UK, while C57BL/6N male mice were obtained from Jackson Laboratory and the colony was bred and maintained in the animal house. Animals were assigned to either a control group (standard chow diet RM1 (P) 801151, Special Diets Services) or the HFpEF group (high-fat diet (Research Diets Inc; D12492i) and L-NAME (Sigma Aldrich, UK; N5751-25G) at concentrations ranging from 0.5 g/L to 1.75 g/L. Mice were kept on HFD+L-NAME between 7 to 15 weeks to induce HFpEF.

For the 3- and 4-hit protocols, 6-week male and female mice were purchased from Animal Bioresources. At 12 weeks of age, mice were randomly assigned to either the control group (standard brown chow, Specialty Feeds Ltd SF00-100) or HFpEF group (high fat diet with 60% energy from lard, Specialty Feeds SF18-072) and were maintained on the diets for time denoted on their relevant protocols (e.g. 6 months for refined protocol or 13 months for full length). After either 6 or 13 months of diet, the HFpEF groups received an intraperitoneal injection of desoxycorticosterone pivalate (75 mg/kg, Zycortal Suspension). One week after DOCP injection, mice started 1% NaCl drinking water (793566, Merck) which continued to the end of the protocol. All experiments were performed in male and female mice, and animals were randomly allocated to procedure groups.

### Exercise testing

For the 2-hit model, mice underwent a three-day acclimatization period for 10-15 min each day on the forced exercise wheel (Lafayette instruments) prior to the intolerance test. During the intolerance test, mice began with a 5-min warm-up at 3 m/min, after which the speed was increased to 4 m/min and maintained until the animals reached exercise intolerance ^13^. Exercise intolerance was defined as the point at which a mouse could not resume running within 10 s after coasting inside the wheel.

For the 4-hit model, mice were acclimated to the treadmill (Ugo Basile) for 15 min on three consecutive days. The treadmill was set at a 20° incline and increased at a rate of 2m/min until the animals were exhausted ^4^. Exhaustion was defined by the animal’s refusal to run after three pushes on the back with a brush. The running time and distance were recorded.

### Blood pressure

Blood pressure was measured non-invasively on conscious mice using the Kent CODA high throughput 6-channel instrument (for 2-hit model, BP-2000 Blood Pressure Analysis System™ (BIOSEBLAB instruments-BP-2000), was used). Mice were acclimated to the restrainers for 10 min per day, for three consecutive days. For testing, the restrainer holding the animal was placed on a 37°C warming tray and covered with a blanket. After allowing 5-10 min for the animals to settle, the occlusion and volume pressure recording cuffs were placed around the tail. 20-25 readings were taken, including 10 acclimatization readings and 10-15 test readings. The measurements were repeated on at least two different days. Animals were not restrained or left in a heated chamber for more than 30 min.

### Echocardiography

Echocardiography was performed using Siemens Acuson Sequoia (for the 2-hit model) and Fujifilm VisualSonics VEVO F2 LAZR-X with a UHF57X transducer (for the 4-hit model). Anaesthesia was induced using 3% isofluorane for three min. Isofluorane was reduced to 1.5-2.5% and adjusted as necessary to maintain the heart rate between 400 to 500 bpm^14^. Left ventricular ejection fraction (LVEF) and global longitudinal strain (GLS) measurements were obtained from a parasternal long axis view, and transmitral doppler measurements and tissue doppler measurements at the mitral annulus were obtained from an apical four chamber view. Images were stored digitally and analyzed blindly using imageJ and VisualSonics workstation.

### Organ weighing

Organs were weighed using an Ohaus Pioneer PX Analytical Balance. Animals were either anesthetized with 4% isoflurane (maintenance: 2%), and reflexes checked before thoracotomy (2-hit model), or pentobarbitol anesthesia used (4-hit model; 100 mg/kg of a 25 mg/ml solution). The heart and lungs were excised, washed with saline, and separated. The aorta was cannulated, and the heart flushed with saline to clear blood, dried and then weighed. Lungs were cleaned of excess tissues and weighed. Lungs were dried using a Speed-Vac Concentrator until weight stabilized after which the wet-to-dry lung weight ratios were calculated for the 2-hit model^4^, or wet lung weight was normalized to tibia length as a marker for lung congestion (4-hit model) ^5^.

## Results

Previously it has been suggested that C57BL/6N mice may be more susceptible to the standard 2-hit model than C57BL/6J animals (Pepin et al, 2023). To better understand the comparative susceptibilities of N and J strains we explored their responses to different treatment regimens that varied both in the dosage and length of treatment of L-NAME (Figure 1 and 2). HFpEF was determined to have been fully induced if there were significant changes in blood pressure, body weight, the E/A ratio, exercise capacity, left ventricular mass, and lung congestion as indicated by increased wet/dry weight ratio.

**Figure. 1.**
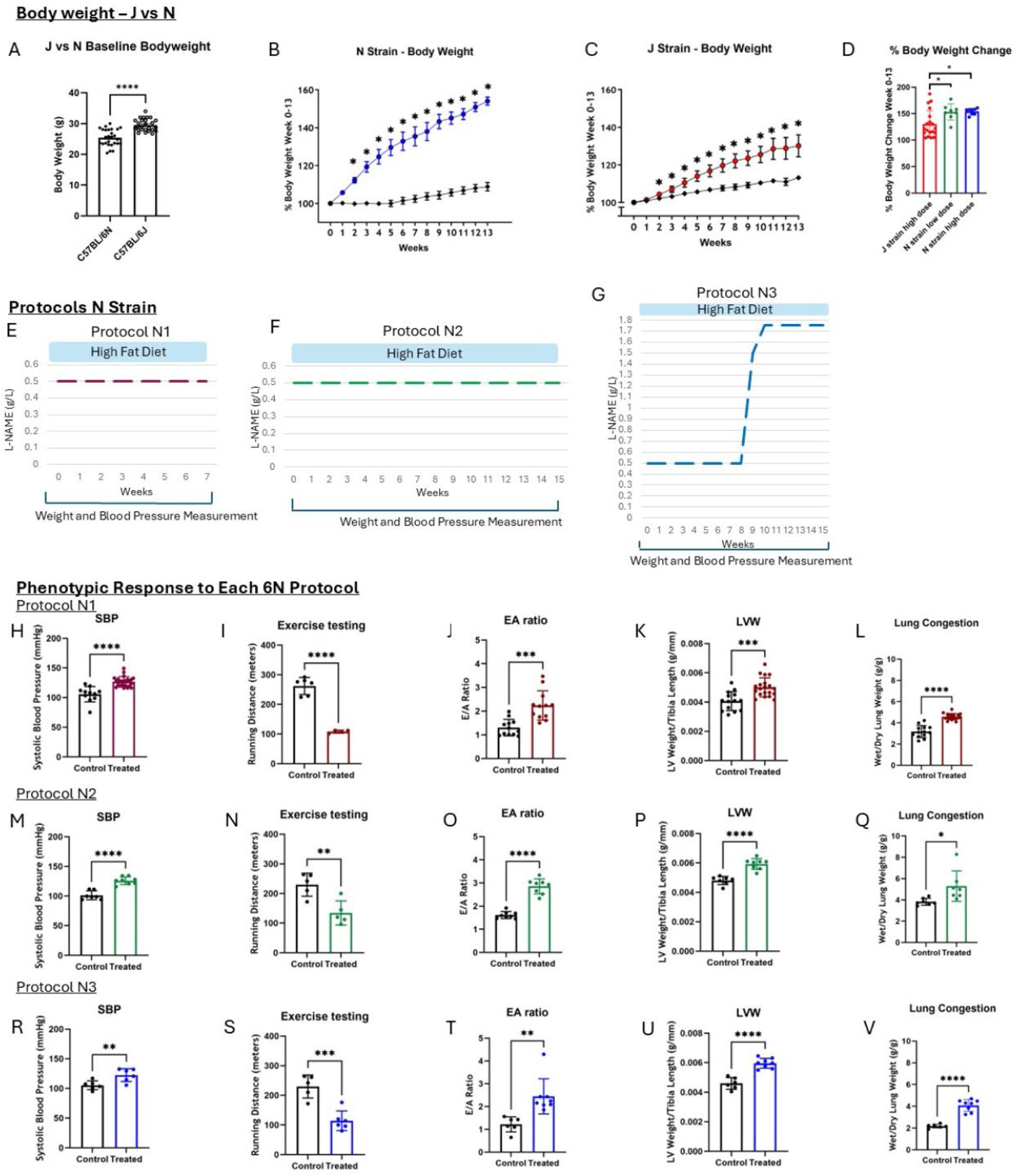
N strain mice produce robust HFpEF phenotype in response to all protocols. A. Baseline body weight of 6J and 6N mice (n = 27 6N mice, n = 26 6J mice). B. Response of N strain body weight to 13 weeks of high fat diet and L-NAME (n = 6 for control, n = 8 for treated). C. Response of J strain body weight to 13 weeks of high fat diet and L-NAME (n = 26 for control, n = 25 for treated). D. Percentage body weight change of J and N strain mice after 13 weeks of high fat diet, and high or low dose L-NAME (n = 19 for J strain high dose, n = 8 for N strain low dose, n = 8 for N strain high dose). E. Schematic of 6N protocol N1, showing 7 weeks of low dose L-NAME. F. Schematic of 6N protocol N2, showing 15 weeks of low dose L-NAME. G. Schematic of 6N protocol N3, showing an increasing dose of L-NAME over 13 weeks. Panels H-K depict measurements taken at the end of the N1 protocol. H. Systolic blood pressure (SBP) of control and treated mice (n = 11 for control, n = 24 for treated). I. Running distance of control and treated mice (n = 6 for control, n = 4 for treated). J. Echocardiography measurement representing peak velocity of mitral blood flow at early filling, to peak velocity of mitral blood flow at late filling (E/A ratio) (n = 12 for control, n = 12 for treated). K. Left ventricular weight (LVW) normalized to tibia length as a measurement of LV hypertrophy (n = 14 for control, n = 20 for treated). L. Ratio of wet to dried lung weight as a measurement of lung congestion (n = 13 for control, n = 18 for treated). Panels M-Q depict measurements taken at the end of the N2 protocol. M. SBP of control and treated mice (n = 7 for control, n = 8 for treated). N. Running distance of control and treated mice (n = 5 for control, n = 5 for treated). O. Echocardiography measurement representing peak velocity of mitral blood flow at early filling, to peak velocity of mitral blood flow at late filling (E/A ratio) (n = 8 for control, n = 8 for treated). P. Left ventricular weight (LVW) normalized to tibia length as a measurement of LV hypertrophy (n = 7 for control, n = 8 for treated). Q. Ratio of wet to dried lung weight as a measurement of lung congestion (n = 7 for control, n = 7 for treated). Panels R-V depict measurements taken at the end of the N3 protocol. R. Systolic blood pressure (SBP) of control and treated mice (n = 6 for control, n = 6 for treated). S. Running distance of control and treated mice (n = 5 for control, n = 6 for treated). T. Echocardiography measurement representing peak velocity of mitral blood flow at early filling, to peak velocity of mitral blood flow at late filling (E/A ratio) (n = 6 for control, n = 8 for treated). U. Left ventricular weight (LVW) normalized to tibia length as a measurement of LV hypertrophy (n = 6 for control, n = 8 for treated). V. Ratio of wet to dried lung weight as a measurement of lung congestion (n = 6 for control, n = 8 for treated). *P<0.05, **P<0.01, ***P<0.001, ****P<0.0001 by unpaired t-test.

**Fig 2.**
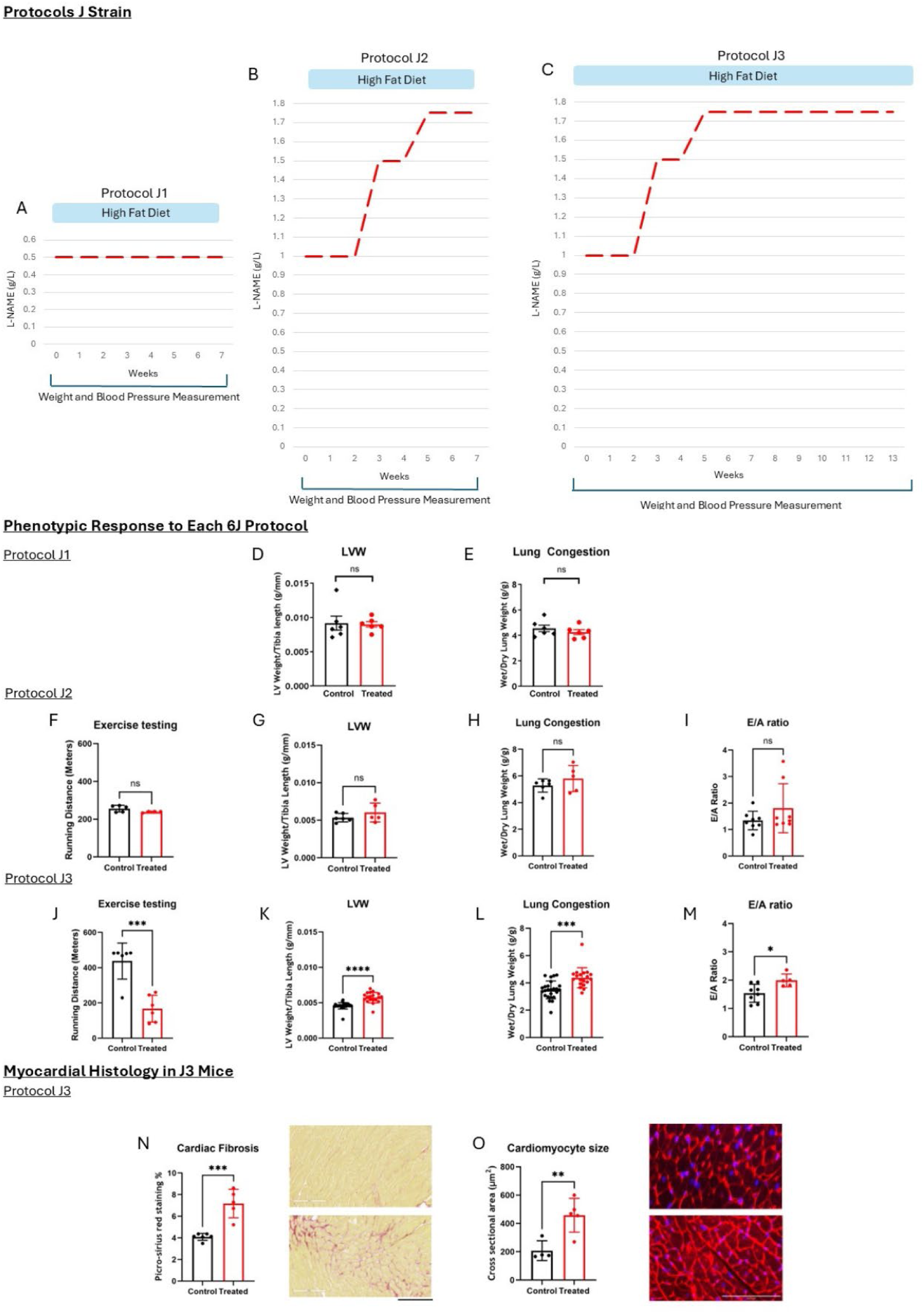
Protocol testing to produce robust HFpEF phenotype in J strain. A. Schematic of 6J protocol J1, showing 7 weeks of low dose L-NAME. B. Schematic of 6J protocol J2, showing an increase of L-NAME over 7 weeks. C. Schematic of 6J protocol J3, showing an increase of L-NAME which is maintained for 13 weeks. D. Left ventricular weight (LVW) normalized to tibia length as a measurement of LV hypertrophy at the end of J1 protocol (n = 6 for control, n = 6 for treated). E. Ratio of wet to dried lung weight as a measurement of lung congestion at the end of J1 protocol (n = 6 for control, n = 6 for treated). Panels F-I show measurements from the end of J2 protocol. F. Running distance of control and treated mice (n = 5 for control, n = 4 for treated). G. LVW normalized to tibia length as a measurement of LV hypertrophy (n = 6 for control, n = 5 for treated). H. Ratio of wet to dried lung weight as a measurement of lung congestion (n = 6 for control, n = 5 for treated). I. Echocardiography measurement representing peak velocity of mitral blood flow at early filling, to peak velocity of mitral blood flow at late filling (E/A ratio) (n = 8 for control, n = 8 for treated). Panels J-M show measurements from the end of J3 protocol. J. Running distance of control and treated mice (n = 6 for control, n = 6 for treated). K. LVW normalized to tibia length as a measurement of LV hypertrophy (n = 26 for control, n = 19 for treated). Ratio of wet to dried lung weight as a measurement of lung congestion (n = 25 for control, n = 19 for treated). N. Increased cardiac fibrosis in J3 mice as shown by picro-sirius red staining (n = 6 for control, n = 5 for treated). O. Cardiomyocyte size in protocol J3 mice (n =4 for control, n = 5 for treated) . *P<0.05, **P<0.01, ***P<0.001, ****P<0.0001 by unpaired t-test.

### Both N and J strain respond to high fat diet and L-NAME

There are contradictory reports in the literature as to whether N- or J-strain mice are intrinsically heavier ^15^. Our results show that age matched J-strain animals had a small but significantly higher baseline body weight than their N-strain counterparts (Figure 1A). In agreement with data reported by Nemoto *et al,* both strains showed significant weight gain on a high fat diet (Figure 1B and C), with percentage weight gain being significantly higher in N-strain animals (Figures 1D) ^15^. These results contrast with others ^12^ who found that mice lacking a functional NNT gene, as is the case in the J-strain, did not show significant weight gain on a high fat diet. Within the more responsive N-strain, animals on a high fat diet showed similar increases in weight using low or high-dose NAME (Figures 1D).

### Higher doses of L-NAME do not augment the HFpEF phenotype in N-strain mice

We next examined whether increasing doses of L-NAME impacted the HFpEF phenotype in N-strain mice. Although length of treatment varied, a single HFD was used throughout (Figure. 1A). We applied three protocols to the N strain mice: HFD + 0.5g/L L-NAME for 7 weeks (Protocol N1, Figure. 1E), HFD + 0.5g/L L-NAME for 15 weeks (Protocol N2, Figure. 1F), or HFD + 0.5g/L L-NAME for 8 weeks, followed by 1.5g/l for two weeks, and then 1.75g/l for 5 weeks (Protocol N3, Figure. 1G).

As expected, ejection fraction did not change in these protocols (Supplemental Figures 1A & B). Although there was some weekly variation, animals treated with 0.5g/l L-NAME showed a rise in systolic blood pressure (SBP) after 1 week of treatment and plateaued thereafter. Continued treatment up to 7 weeks (Protocol N1, Figure. 1H) led to significantly increased blood pressure, reduction in running distance (Figure. 1I), increase in E/A ratio (Figure. 1J), increase in left ventricular mass (Figure. 1K), and increase in wet/dry lung weight (Figure. 1L). The N2 (Figure. 1M-Q) and N3 (Figure. 1Q-U) protocols also led to significant changes in these parameters, but no more markedly than the N1 protocol. These results demonstrate that in N-strain animals a robust HFpEF phenotype can be achieved after 7 weeks at an L-NAME dose of 0.5g/l and that higher doses do not augment the phenotype further.

### HFpEF can be induced in J-strain mice

Producing a robust phenotype in the J-strain would facilitate research using genetic mice models which are mostly based on the J-strain. Given previous reports of resistance to HFpEF in the J-strain (Pepin et al., 2023) we tested different protocols, varying the length of treatment and dose of L-NAME, to determine whether a robust phenotype is possible in the J-strain. As protracted 15-week L-NAME treatment at 0.5 g/l did not augment blood pressure, we changed to a ramped dose protocol as per below.

J-strain mice treated for 7 weeks at doses of L-NAME ranging from 0.5-1.75 g/l (Protocols J1, Figure. 2A; J2, Figure. 2B) failed to show the full range of signs indicative of HFpEF (Figure 2D-I). This contrasts with the N-strain where 7 weeks of HFD and 0.5g/L of L-NAME could achieve a comprehensive HFpEF phenotype. However, J strain mice treated for 13 weeks with a dosage regime rising from 1.0g/l to 1.75g/l (Protocol J3, Figure. 2C) showed significant changes in exercise intolerance, left ventricular mass, and lung wet/dry ratio and E/A ratio (Figure 2 J-M). In addition, postmortem histological examination revealed increased cardiac fibrosis and increased cardiomyocyte cross sectional area indicating cardiac hypertrophy (Figures 2N &O). Importantly the ejection fraction was preserved in these animals (Supplemental Figure 1C & D). These results clearly demonstrate that it is possible to induce a HFpEF phenotype in C57BL/6J mice, in which we also confirmed histologically the presence of cardiomyocyte hypertrophy and extracellular myocardial fibrosis.

### Augmenting the phenotype in 3-Hit mice: The “4-Hit” protocol

The 3-hit model described by Deng et al in 2021 is schematically outlined alongside our modification with NaCl (“4-Hit’) in Figure. 3A. Using the 3-hit model, we did not see a significant elevation of SBP at 13 months in male C57BL/6N mice, whereas we did after 1% NaCl was added to drinking water (Figure. 3B). We saw a similar pattern for DBP (Figure. 3C), E/e’ ratio (Figure. 3D), and GLS (Figure. 3E). Left ventricular ejection fraction was preserved in both protocols (Figure. 3F), and mice fed high fat diet were significantly heavier (Figure. 3G). Due to the observed enhancement of the HFpEF phenotype with the addition of 1% NaCl drinking water, it was provided 1 week post DOCP injection in subsequent batches and referred to as 4-hit model. Hypertension was sustained for approximately 7 weeks before decreasing (Supplemental Figure 1E-F), confirming the NaCl alone was not inducing the hypertensive effect, and we also determined that the mice could tolerate a second dose of DOCP with no adverse response, allowing the HFpEF phenotype to be sustained longer. Importantly, ejection fraction was maintained in all cohorts (Supplementary Figures 1 G-J). Further, we demonstrated we could recreate the HFpEF phenotype with the 4-hit protocol in a shorter timeframe, as demonstrated in the 6J 4-hit mice in which the initial high fat diet loading phase was shortened by 5 months (Figure. 4G-L, 4S-X, Schematic in Supplementary Figure 2A).

**Figure. 3.**
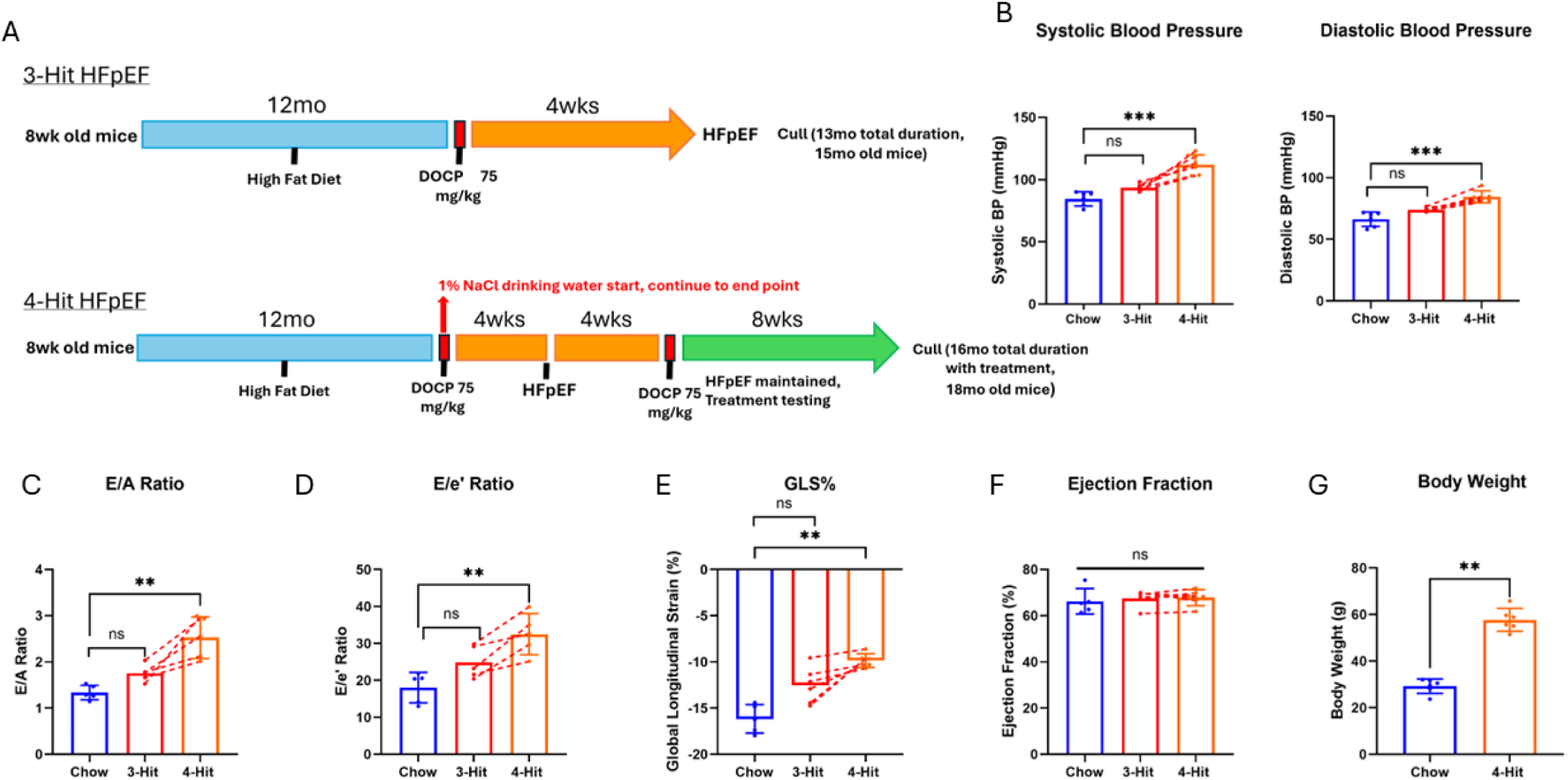
A reproducible HFpEF phenotype incorporating aging – “4-Hit”. A. Schematic depicting the traditional 3-Hit approach, and our modified 4-hit approach incorporating NaCl drinking water and additional DOCP injection. B. Systolic and diastolic blood pressure after the traditional 3-Hit approach in red, and following the addition of NaCl in orange. C. Echocardiography measurement representing peak velocity of mitral blood flow at early filling, to peak velocity of mitral blood flow at late filling (E/A ratio). D. Ratio of peak velocity of mitral blood flow at early filling to peak early diastolic mitral annulus velocity (E/e’) before and after the addition of NaCl. E. Global longitudinal strain (GLS) of the left ventricle. F. Preserved ejection fraction of the left ventricle. G. Body weight of healthy chow mice and 4-Hit mice at sacrifice. For all panels male C57BL/6N mice were used, n = 6 for chow, n = 7 for 3-Hit, n = 7 for 4-Hit. *P<0.05, **P<0.01, ***P<0.001, ****P<0.0001 by Mann-Whitney test.

**Figure. 4.**
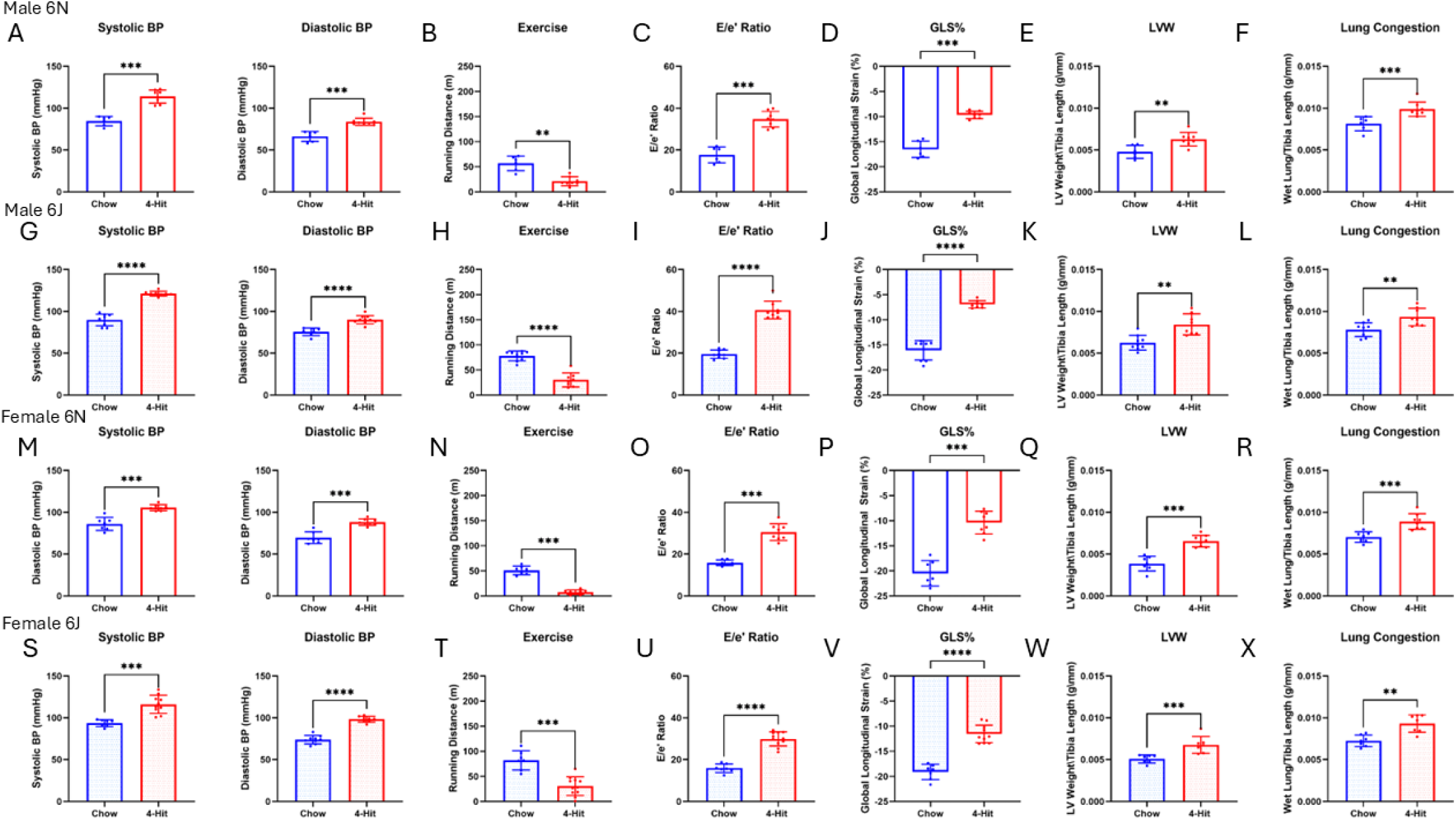
4-Hit model reproducibly produces HFpEF in male and female J and N. Panels A-F depict measurements from male 6N mice at the end of 4-Hit protocol. A. Systolic and diastolic blood pressure of chow and 4-Hit male 6N mice. B. Running distance of chow and 4-Hit mice. C. Ratio of peak velocity of mitral blood flow at early filling to peak early diastolic mitral annulus velocity (E/e’). D. Global longitudinal strain (GLS) of the left ventricle. E. Left ventricular weight (LVW) normalized to tibia length as a measurement of LV hypertrophy. F. Wet lung weight normalized to tibia length as a measurement of lung congestion. For A-F n = 7 for chow, n = 8 for 4-hit. Panels G-L depict measurements from male 6J mice at the end of 4-Hit protocol. G. Systolic and diastolic blood pressure of chow and 4-Hit male 6J mice. H. Running distance of chow and 4-Hit mice. I. Ratio of peak velocity of mitral blood flow at early filling to peak early diastolic mitral annulus velocity (E/e’). J. Global longitudinal strain (GLS) of the left ventricle. K. LVW normalized to tibia length as a measurement of LV hypertrophy. L. Wet lung weight normalized to tibia length as a measurement of lung congestion. For G-L n = 9 for chow, n = 8 for 4-Hit. Panels M-R depict measurements from female 6N mice at the end of 4-Hit protocol. M. Systolic and diastolic blood pressure of chow and 4-Hit female 6N mice. N. Running distance of chow and 4-Hit mice. O. Ratio of peak velocity of mitral blood flow at early filling to peak early diastolic mitral annulus velocity (E/e’). P. Global longitudinal strain (GLS) of the left ventricle. Q. LVW normalized to tibia length as a measurement of LV hypertrophy. R. Wet lung weight normalized to tibia length as a measurement of lung congestion. For M-R n = 7 for chow, n = 8 for 4-Hit. Panels S-X depict measurements from female 6J mice at the end of 4-Hit protocol. S. Systolic and diastolic blood pressure of chow and 4-Hit female 6J mice. T. Running distance of chow and 4-Hit mice. U. Ratio of peak velocity of mitral blood flow at early filling to peak early diastolic mitral annulus velocity (E/e’). V. Global longitudinal strain (GLS) of the left ventricle. W. LVW normalized to tibia length as a measurement of LV hypertrophy. X. Wet lung weight normalized to tibia length as a measurement of lung congestion. For S-X n = 7 for chow, n = 10 for 4-Hit. *P<0.05, **P<0.01, ***P<0.001, ****P<0.0001 by Mann-Whitney test.

To ensure the aged C57BL/6J mice still responded to the DOCP and NaCl in the expected manner, a group of 6J male mice were aged to 13 months, injected with DOCP and given NaCl drinking water. We observed the same increase in systolic and diastolic blood pressure (Supplementary Figure 2B), and as inducing sustained hypertension is the main limiting factor in aged HFpEF models, there are no indications our findings would not translate if the protocol was not shortened by 5 months in this strain.

We next proceed to examine the key HFpEF parameters in the 4-Hit model across substrain and sex (Figure. 4). SBP was significantly elevated in the 4-Hit model in male 6N mice, as was DBP (Figure. 4A), with significant reduction in exercise tolerance (Figure. 4B), increase in E/e’ ratio (Figure. 4C), significant decrease in GLS (Figure. 4D), and significantly increased LV mass (Figure. 4E), and significantly increased wet/dry lung weight (Figure. 4F). The same pattern, with even greater significance, was seen in Male 6J mice (Figure. 4G-L). A similar pattern was observed in female 6N mice (Figure. 4M-R), albeit with slightly less significance. Similar to the male mice, in female 6J the significance of the changes was somewhat greater (Figure. 4S-X).

### Phenotype comparison: 4-Hit vs 2-Hit

We next compared the 4-Hit to the traditional “standard” 2-Hit HFpEF phenotypic parameters in male and female J-strain mice. These 2-Hit mice followed the protocol as outlined in Figure 1F (originally described by Schiattarella *et al*^4^.) though they are C57BL/6J background not 6N. In the Male 6J mice, significance was greater across all components of the phenotype (Figure. 5A-F) compared to the 2-Hit model (Figure. 5G-L). However, in female mice, if one considers the presence of all parameters necessary for a successful mode, then the 4-Hit model (Figure. 5M-R) was clearly superior to the standard 2-Hit model (Figure. 5S-X) in female 6J mice. Specifically, the left ventricular mass and lung congestion were not significant in the standard 2-Hit model either male or female 6J mice. Interestingly, in both male and female 6J mice the 2-Hit model was less robust in producing exercise intolerance compared to the 4-Hit model. Lung congestion is a crucial parameter of HFpEF and was not observed in these 2-Hit mice using the unmodified protocol, highlighting the importance of the modifications to the model outlined in this paper.

**Figure. 5.**
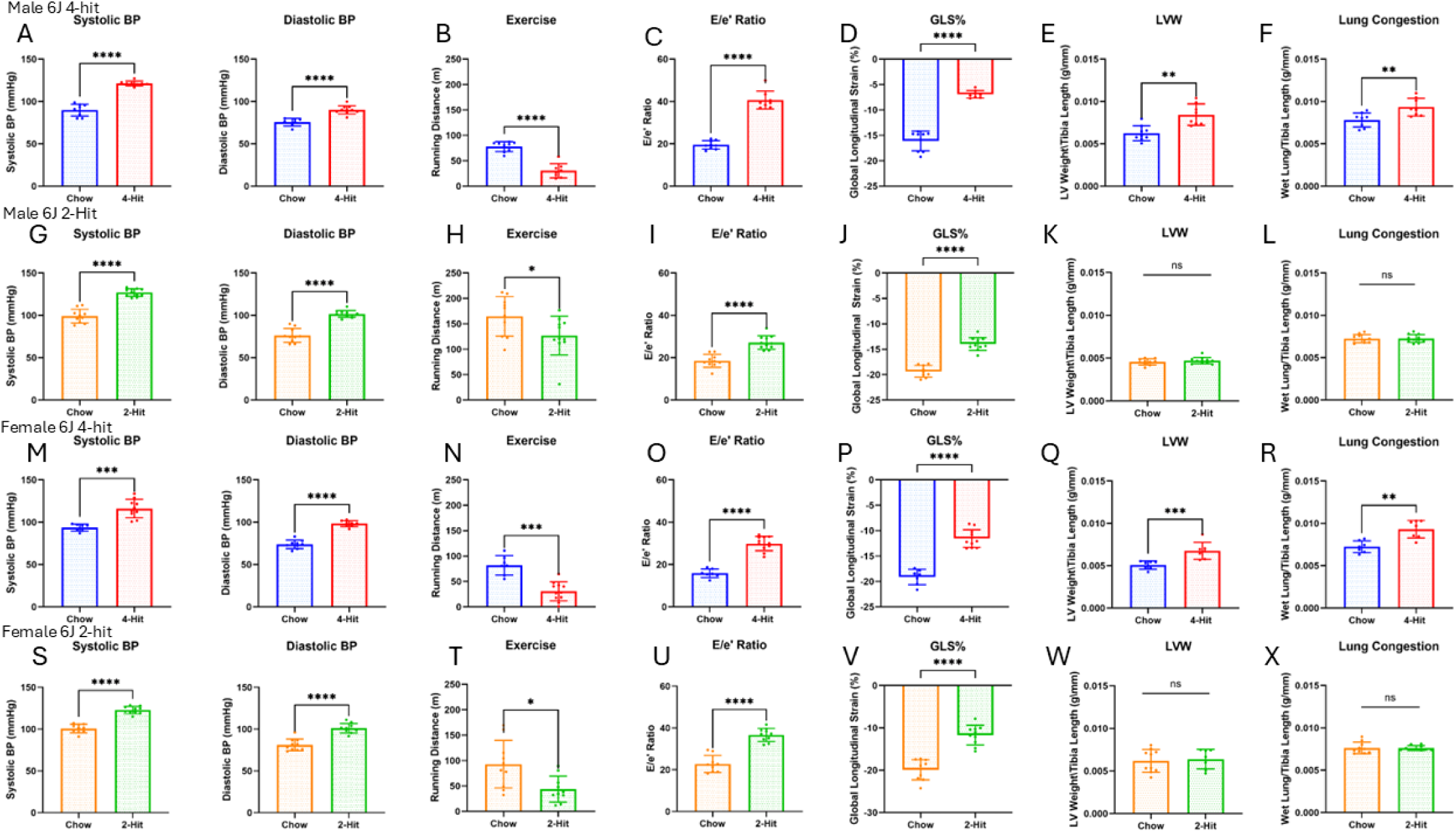
Phenotypic comparison of 4-Hit (incorporating aging) vs standard 2-Hit (no aging). 4-Hit data is duplicated from figure 4. Panels A-F depict measurements from 6J males at completion of 4-Hit protocol. A. Systolic and diastolic blood pressure of chow and 4-Hit male 6J mice. B. Running distance of chow and 4-Hit mice. C. Ratio of peak velocity of mitral blood flow at early filling to peak early diastolic mitral annulus velocity (E/e’). D. Global longitudinal strain (GLS) of the left ventricle. E. Left ventricular weight (LVW) normalized to tibia length as a measurement of LV hypertrophy. F. Wet lung weight normalized to tibia length as a measurement of lung congestion. For A-F, n = 7 for chow, n = 8 for 4-Hit. Panels G-L depict measurements from 6J males at completion of the standard 2-Hit protocol. G. Systolic and diastolic blood pressure of chow and 2-Hit male 6J mice. H. Running distance of chow and 2-Hit mice. I. Ratio of peak velocity of mitral blood flow at early filling to peak early diastolic mitral annulus velocity (E/e’). J. Global longitudinal strain (GLS) of the left ventricle. K. LVW normalized to tibia length as a measurement of LV hypertrophy. L. Wet lung weight normalized to tibia length as a measurement of lung congestion. For G-L, n = 10 for chow, n = 11 for 2-Hit. Panels M-R depict measurements from 6J females at completion of 4-Hit protocol. M. Systolic and diastolic blood pressure of chow and 4-Hit female 6J mice. N. Running distance of chow and 4-Hit mice. O. Ratio of peak velocity of mitral blood flow at early filling to peak early diastolic mitral annulus velocity (E/e’). P. Global longitudinal strain (GLS) of the left ventricle. Q. LVW normalized to tibia length as a measurement of LV hypertrophy. R. Wet lung weight normalized to tibia length as a measurement of lung congestion. For M-R, n = 7 for chow, n = 10 for 4-Hit. Panels S-X depict measurements from 6J females at completion of the standard 2-Hit protocol. S. Systolic and diastolic blood pressure of chow and 2-Hit female 6J mice. T. Running distance of chow and 2-Hit mice. U. Ratio of peak velocity of mitral blood flow at early filling to peak early diastolic mitral annulus velocity (E/e’). V. Global longitudinal strain (GLS) of the left ventricle. W. LVW normalized to tibia length as a measurement of LV hypertrophy. X. Wet lung weight normalized to tibia length as a measurement of lung congestion. For S-X n = 10 for chow, n = 10 for 2-Hit. *P<0.05, **P<0.01, ***P<0.001, ****P<0.0001 by Mann-Whitney test.

## Discussion

While murine models play a very important role in the investigation of HFpEF, capturing a more complete clinical picture of the disease state with such complex pathophysiology is not without limitations. The aetiology of insidious onset cardiac disease, such as HFpEF, is complex and derives from multiple, interlinked predisposing risk factors such as age, hypertension, type II diabetes, obesity, kidney disease, COPD, anaemia and sleep disordered breathing ^16,17^. This complex causation is paralleled by the complexity of the different pathophysiological changes associated with the clinical presentation of HFpEF including cardiac fibrosis and stiffness, metabolic changes, mitochondrial and energy dysfunction, microvascular disease, and inflammatory changes, all of which contribute to reduced ventricle compliance, suboptimal diastolic filling, and increased filling pressures ^2,18–20^.

If mouse models are to be useful, the minimum requirement is that they are robust, and the results are highly repeatable. Only if these conditions are satisfied can researchers rely on the model to perform as expected and use it as a foundation to build our knowledge of the condition under study. However, a potential pitfall relates to the phenotypic differences between strains and sub-strains, and insufficient attention to this can lead to discrepancies. The strains investigated here genetically diverged in the 1950s when separate breeding colonies were maintained in the Jacks Laboratory and NIH labs, hence the J- and N-strains and have been isolated for more than 200 generations ^21^. Despite their common origin, several studies have emphasised the numerous genomic changes including structural variations, indel and single nucleotide polymorphisms (SNPs) that distinguish the two strains ^21^ resulting in a large number of identified phenotypic differences ^22^ including fundamental differences in cardiac physiology. C57BL/6N are often used for the development of heart failure models and demonstrate changes in connective tissue markers Nppa, and Myh7, and relatively more LV fibrosis following 14 days of angiotensin II infusion^23^. Additionally, Zi *et al*^24^ determined that C57BL/6N mice developed eccentric hypertrophy with cardiac deterioration depending on age and time course, where as C57BL/6J mice developed variable cardiac phenotypes, when both strains underwent the same exposure to transverse aortic constriction (TAC).

In 2019 Schiattarella *et al* ^4^ reported a new rodent model for HFpEF. Using the remarkably simple approach of induced obesity combined with high systolic blood pressure they were able to produce a range of signs of consistent with HFpEF in a matter of weeks in N-strain mice. Subsequently Pepin et al ^12^ discussed the repeatability of this approach and the difficultly of reproducing this phenotype in other mouse strains. They showed that resistance to the induction of HFpEF segregated with a mutated form of the NNT gene that was harboured by the J-strain. However, several phenotypic characteristics, including many of them relevant to HFpEF, such as energy metabolism, glucose tolerance, body weight and response to cardiac insult could not be explained by alternations in the NNT gene alone ^15,25^.

Many genetically modified mouse models are generated in C57BL/6J strains and as such they occupy a prominent place in model-based research. It was important, therefore, to investigate if it was possible to induce the signs of HFpEF in the resistant J-strain. In this report, we demonstrate that by extending the treatment period and utilising a higher dose of the hypertension-inducing drug L-NAME it was possible to produce the panoply of changes that represent many of the signs associated with human heart failure with preserved ejection fraction.

The literature contains mixed reports as a number of groups ^9,26,27^ have successfully utilised J-strain mice to induce HFpEF at low doses of L-NAME and relatively short treatment periods of 5-8 weeks. These results are in stark contrast to the results reported here and previous reports (Pepin et al, 2023).

An additional complexity surrounding HFpEF murine models is incorporating the ageing component, a prominent risk factor in the human HFpEF patient that cannot be overlooked. As NOS activity, the target of L-NAME-induced hypertension, decreases with age ^28^, the L-NAME model is less effective in the aged rodent.

Recognizing these limitations, Deng et al developed a “3-hit” model, incorporating aging as an additional factor to account for the increased risk of HFpEF in older populations (Deng, et al., 2021). In this approach, 3-month-old mice are fed a high-fat diet for 13 months to induce metabolic stress, and in the final month, desoxycorticosterone pivalate (DOCP) is administered intraperitoneally to induce hypertension and systemic inflammation. This method has been regarded as the gold standard for HFpEF models, as it successfully integrates metabolic stress, hypertension, and aging—three key drivers of HFpEF, making it more representative of the human condition ^6,7^.

DOCP, a long-acting synthetic mineralocorticoid, mimics the effects of aldosterone by binding to mineralocorticoid receptors in the kidney’s distal tubules, promoting sodium retention and potassium excretion. This increase in sodium retention leads to an expansion of extracellular fluid volume, which can elevate blood pressure. However, under normal physiological conditions, the body activates compensatory mechanisms, particularly the Renin-Angiotensin-Aldosterone System (RAAS), to restore sodium balance and limit further increases in blood pressure ^29,30^.

While exogenous DOCP administration leads to an initial retention of sodium, this effect is often transient. Over time, the phenomenon of ’aldosterone escape’ occurs, wherein the kidneys adjust sodium excretion to match intake despite the continued presence of aldosterone ^31^. This adjustment helps prevent further fluid retention, although blood pressure may remain elevated. Thus, sodium retention alone may not be sufficient to sustain prolonged hypertension.

To overcome these compensatory mechanisms, our model combines DOCP with additional salt supplementation. While DOCP promotes sodium retention, the supplemental sodium load ensures that the kidneys are persistently challenged, preventing compensatory RAAS-driven sodium excretion and sustaining a positive sodium balance, leading to prolonged hypertension. By employing this strategy, we aim to prevent aldosterone escape, sustain hypertension, and promote fluid retention. Therefore, to address the limitations of the 3-hit model and ensure the full HFpEF phenotype, we developed a “4-hit” HFpEF model by adding 1% sodium chloride in the drinking water as the fourth stressor. Crucially, this model worked in C57BL/6N and C57BL/6J strains, and in both male and female mice, also allowing its application in genetic settings.

When working with mouse models, animal welfare considerations should always be at the forefront, and an important modification to the 4-Hit model was determining that we could recreate the HFpEF phenotype with the 4-Hit protocol in a shorter timeframe. Studies have shown very long-term high fat diet exposure to mice can result in detrimental gut issues, dental abnormalities, and exacerbation of genetic predisposition to dermatitis ^32–34^. Furthermore, the desired metabolic effects of the high fat diet, such as weight gain and disturbance of glucose metabolism, are observed in as short as 7 weeks on the 2-Hit L-NAME protocol, so 13 months may appear unnecessary and expensive.

We also confirmed the same phenotype was captured when C57BL/6N mice were aged on healthy brown chow diet for 5 months, then swapped to high fat diet for 6 months (supplemental figure 2 C-O). This resulted in the same age at sacrifice as those on high fat diet for 13 months, but observed no additional phenotypic benefit, and with less unwanted side effects and suffering for the rodents.

## Conclusions

In this paper, we investigated the development of the two-hit model of HFpEF in both C57BL/6N and C57BL/6J strains. Contrary to previous reports, we demonstrated that it is possible to induce HFpEF in J-strain mice by intensifying the traditional two-hit approach. This finding allows researchers to use genetically engineered mouse models in either strain to investigate cardiac and diastolic dysfunction.

Furthermore, we provided a novel robust, reproducible 4-hit aging HFpEF model, which allows for greater clinical translation of results, as the 4-Hit HFpEF model recapitulates the critical risk factors in the human HFpEF patient. We demonstrated this model to work in C57BL/6J and 6N male and female mice, and offered refinements to the protocol which still produce a strong phenotype with animal welfare and protocol timelines in mind. We also demonstrated that the 4-hit model can sustain the HFpEF phenotype for up to 12 weeks, enabling therapeutic options to be tested in this setting.

## Limitations

Although the models herein are the most commonly used and critically appraised murine HFpEF models, they are not the only HFpEF models in use, and other aspects of HFpEF such as chronic pulmonary and renal disease are not incorporated.

## Acknowledgments

We thank Sydney Imaging Preclinical Facility for access to the cardiac imaging instruments. The authors thank Michael Dunne, Margaret Bell, and Catherine Hawksby at the University of Glasgow for their technical assistance.

