## Supplemental Figures for "Optimized protocols for commonly-used murine models of heart failure with preserved ejection fraction"

### Supplemental Figure 1

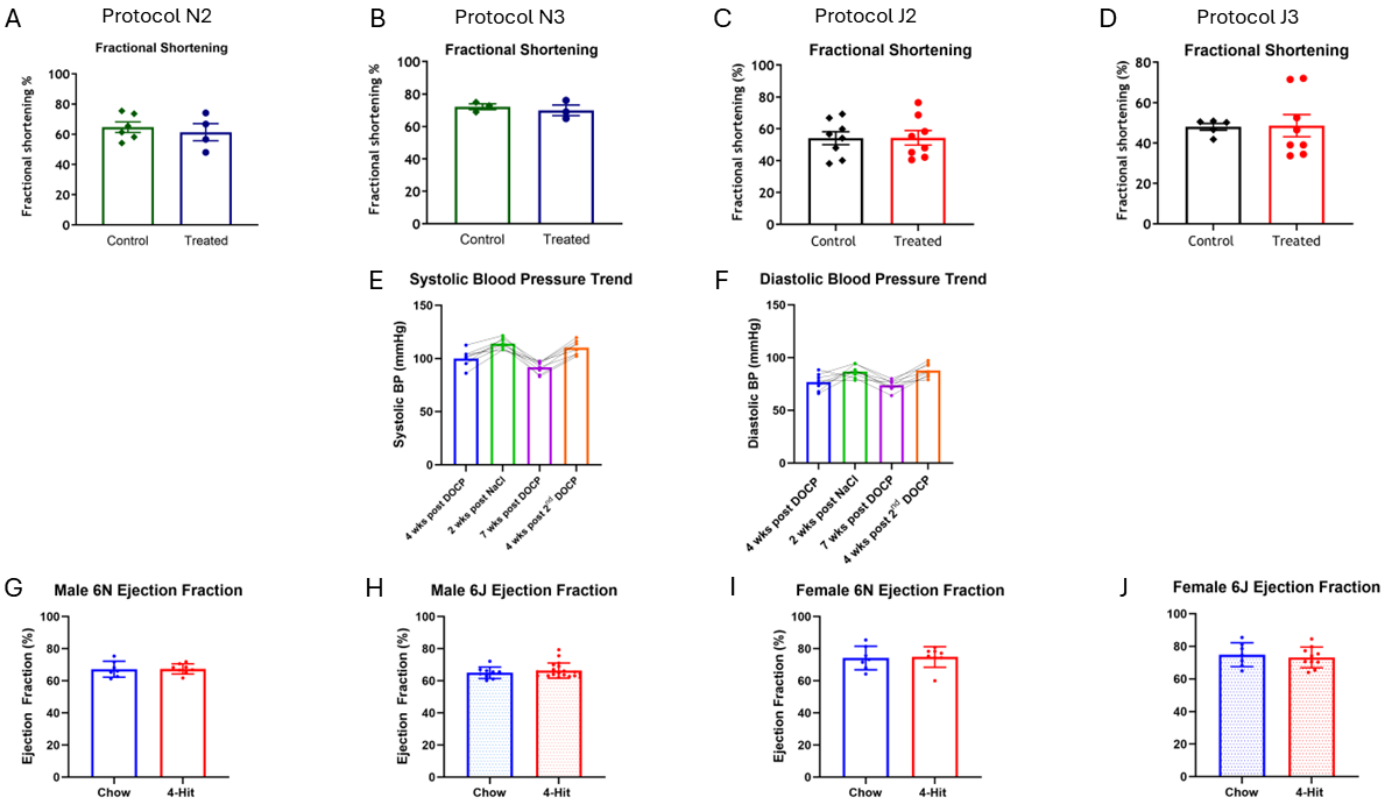


**Supplemental Figure 1. Maintenance of ejection fraction and blood pressure. A. Fractional shortening maintained in mice at conclusion of protocol N2 (n = 6 for control, n = 4 for treated). B. Fractional shortening maintained in mice at conclusion of protocol N3 (n = 3 for control, n = 3 for treated). C. Fractional shortening maintained in mice at conclusion of protocol J2 (n = 8 for control, n = 8 for treated). D. fractional shortening maintained in mice at conclusion of J3 protocol (n = 5 for control, n = 8 for treated). E. Trend of systolic blood pressure in 6N mice between DOCP dosing and addition of NaCl (n = 7). F. Trend of diastolic blood pressure in 6N mice between DOCP dosing and addition of NaCl. G. Ejection fraction (EF) of male 6N mice at conclusion of 4-Hit protocol (n = 7 for chow, n = 8 for 4-Hit). H. EF of male 6J mice at conclusion of 4-Hit protocol (n = 9 for chow, n = 19 for 4-Hit). I. EF of female 6N mice at conclusion of 4-Hit protocol (n = 7 for chow, n = 8 for 4-Hit). J. EF of female 6J mice at conclusion of 4-Hit protocol (n = 7 for chow, n = 10 for 4-Hit).**

### Supplementary Figure 2

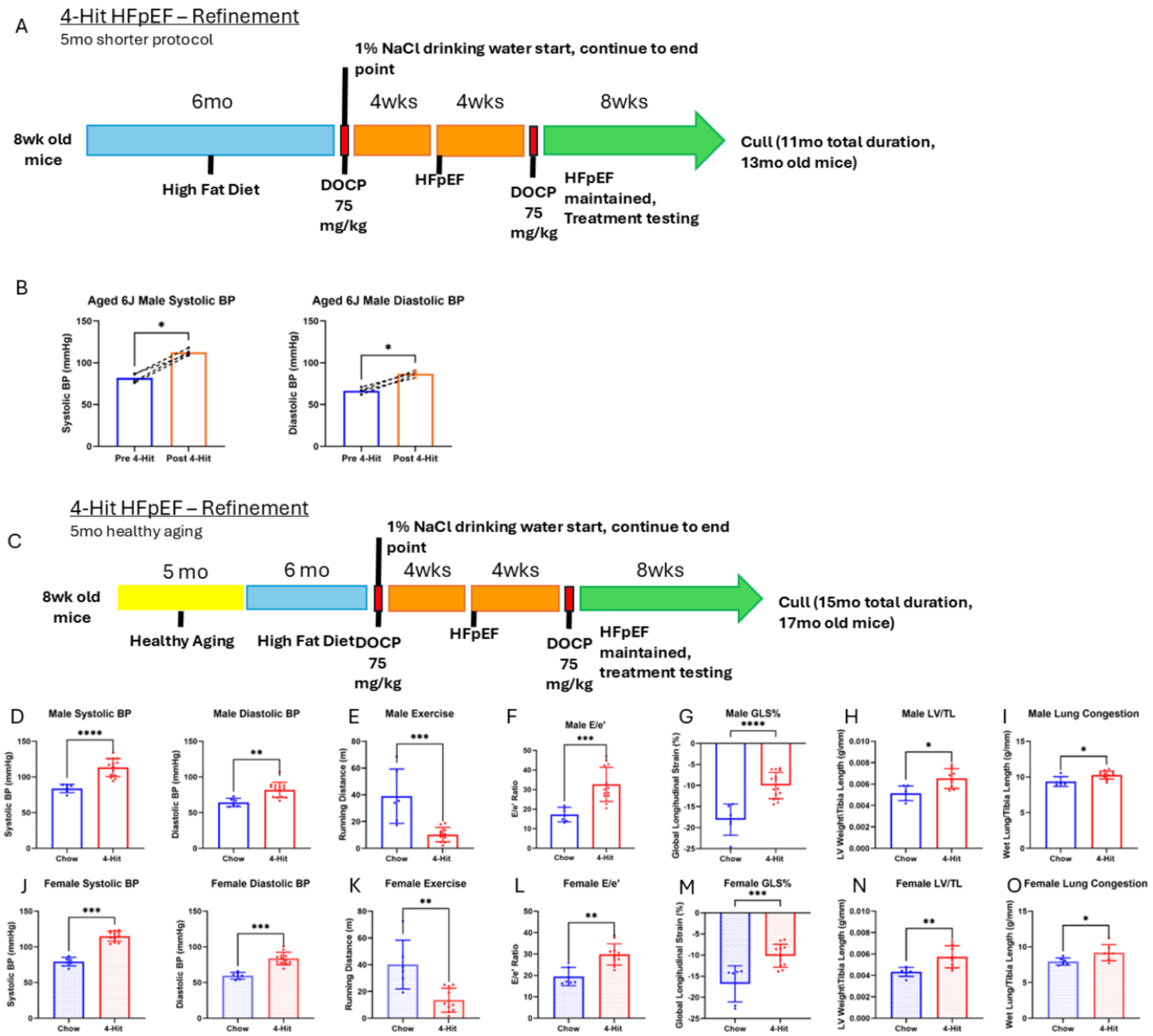


**Supplementary figure 2. Refinements to the 4-Hit model for improved animal welfare. A. Schematic of a refinement option for 4-Hit model with high fat diet (HFD) loading phase shortened by 5 months, for overall duration of 11 months. B. Systolic and diastolic blood pressure of male 6J mice after full duration of 13 months aging, DOCP injection and NaCl drinking water (n = 4). C. A schematic for refinement option for 4-Hit model including 5 months of healthy aging then 6 months high fat diet, for 15 months total duration. Panels D-O depict measurements from male and female 6J mice after completion of the refined 4-Hit model outlined in panel C. D. Systolic and diastolic blood pressure of male mice. E. Running distance of male mice. F. Ratio of peak velocity of mitral blood flow at early filling**

**to peak early diastolic mitral annulus velocity (E/e'). G. Global longitudinal strain (GLS) of the left ventricle. H. Left ventricular weight to tibia length as an indicator of left ventricular hypertrophy. I. Wet lung weight to tibia length as an indicator of lung congestion. For D-I, n = 6 for chow, n = 14 for 4-Hit. J. Systolic and diastolic blood pressure of female mice. K. Running distance of female mice. L. Ratio of peak velocity of mitral blood flow at early filling to peak early diastolic mitral annulus velocity (E/e'). M. Global longitudinal strain (GLS) of the left ventricle. N. Left ventricular weight to tibia length as an indicator of left ventricular hypertrophy. O. Wet lung weight to tibia length as an indicator of lung congestion. For J-O, n = 6 for chow, n = 10 for 4-Hit. *P<0.05, **P<0.01, ***P<0.001, ****P<0.0001 by Mann-Whitney test.**
